# Teaching others reveals hidden structure in conceptual cognitive maps

**DOI:** 10.64898/2026.08.04.742691

**Authors:** Xiaoyan Wu, Peizheng Wu, Wolfgang Omlor, Wolfram Hinzen, Iris E. Sommer, Philipp Homan

**Affiliations:** Department of Adult Psychiatry and Psychotherapy, University of Zurich, Zurich, Switzerland; Neuroscience Center Zurich, University of Zurich and ETH Zurich, Zurich, Switzerland; Department of Translation and Language Sciences, Universitat Pompeu Fabra, Barcelona, Spain; Institució Catalana de Recerca i Estudis Avançats (ICREA), Barcelona, Spain; Department of Neuroscience, University Medical Center Groningen, Groningen, the Netherlands

## Abstract

Disorganized thought is a core feature of the psychosis spectrum. It is recognized, clinically and in daily life, through language, yet its proposed cognitive basis, shallow cognitive maps, the relational structures that organize knowledge, has been tested almost exclusively with task performance and neural measures. Here we show that the organization of a learned conceptual map can be read out from natural speech. Adults varying in schizotypal cognitive disorganization (*n* = 107) learned the same two-dimensional conceptual structure from images or sentences, then described their memory strategy and explained the material to a novice. Explaining elicited two-dimensional spatial descriptions selectively after image-based learning. Higher cognitive disorganization was associated with answers that tracked the question asked less closely, on an automated, rater-free embedding measure robust to report language, again selectively after image-based learning. Natural speech thus provides a scalable behavioral readout of conceptual map organization and its disorganization.

## Introduction

Disorganized thought is among the most striking features of the psychosis spectrum, and it is recognized, in the clinic and in daily life, through language. Its cognitive basis nevertheless remains debated. One influential proposal holds that it reflects shallow or unstable cognitive maps: the relational structures that place items in a mental space and support inference and generalization^1,2^. Cognitive maps were first characterized in spatial navigation^3–5^, but comparable relational codes organize abstract conceptual knowledge across nonspatial domains^6–9^. Evidence for shallow maps in psychosis, however, comes almost entirely from task performance and neural measures, leaving open whether disrupted map structure is expressed in the very behavior through which disorganization is recognized: communicative speech.

This gap matters because a speech-based readout would connect the mechanistic account to the clinical phenomenon itself. Recent work shows that naturalistic speech carries information about the structure of semantic representations and about its disruption in psychiatric conditions^10,11^. Whether the relational structure of a conceptual map can be read out from speech is unknown, in part because people rarely describe relational structure spontaneously. A manipulation is therefore needed that draws latent structure into the open. Explanation is a natural candidate: when people explain what they know to someone else, they reorganize information into relational formats that support communication and understanding^12^, and may thereby externalize structure that remains implicit during self-reflection.

A second consideration concerns how a map is acquired. Language supplies relational structure explicitly, whereas visual input rarely labels relations and requires learners to derive them internally. We recently found that cognitive maps built from visual, but not linguistic, input are selectively disrupted by schizotypy-related cognitive disorganization (CD)^13^, indicating that individual differences in relational organization become visible specifically when structure must be actively constructed. Here we bring these ideas together. Participants learned the same two-dimensional conceptual structure from either images or sentences, then reported their own memory strategy and explained the material to a novice. We asked whether explanation externalizes spatial structure in speech, whether this depends on visual learning, and whether CD attenuates it. Speech was assessed with two independent readouts: human coding of the strategies participants described, and a rater-free measure of how closely their answers tracked the question they had been asked, derived from sentence embeddings.

## Results

Participants (*n* = 107) learned associations between two conceptual dimensions (age group and time of day), forming a two-dimensional (2D) conceptual structure (Figure 1A). During learning, participants completed ordering, association, and navigation tasks designed to promote relational knowledge of this conceptual space. The same structure was presented either image-based or text-based (Figure 1B).

**Figure 1.**
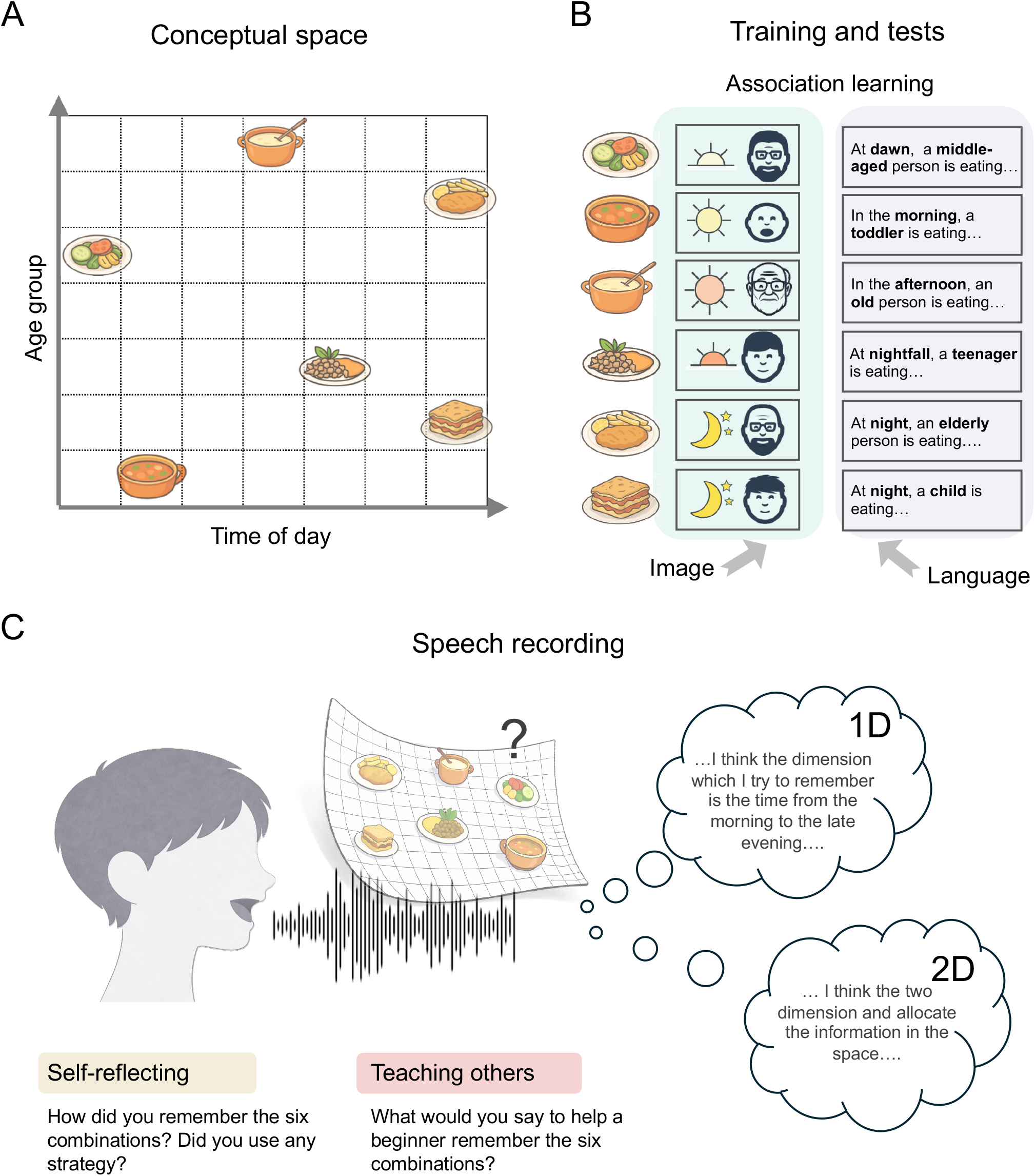
Experimental paradigm and analysis of verbal reports. A.Conceptual space defined by age group (vertical axis) and time of day (horizontal axis). Each food item corresponded to a unique combination of the two conceptual dimensions. **B**. Training phase. Participants learned the food–attribute associations either through images (image-based condition) or linguistic descriptions (language-based condition). **C**. Verbal reports. After learning, participants described either how they remembered the combinations (strategy reflection) or how they would explain them to someone else (teaching explanation). Reports were coded as 1D, 2D, or no organizing strategy.

After learning, participants produced two types of verbal reports: (i) reflecting on their own memory strategies and (ii) explaining how they would teach the information to someone else. Reports were coded as describing a sequential (one-dimensional, 1D), spatial (2D), or no organizing strategy (Cohen’s *κ* = 0.85; Figure 1C).

### Explanation externalizes spatial structure after visual learning

We first tested whether explaining a strategy to others promotes spatial organization in speech. Compared with self-reflection, participants were more likely to describe 2D spatial strategies when teaching (*β* = 1.09, 95% CI [0.24, 1.93], *p* = .012; Supplementary Table 4). This effect was driven by the image-based condition, in which 2D descriptions increased from self-reflection to teaching (exact McNemar test, *p* = .003; Figure 2A) while reports of no strategy decreased (*p* = .003), indicating a shift from unstructured to spatially structured accounts rather than a general increase in verbosity. No such shift occurred after language-based learning (exact McNemar test, *p* = .688; Figure 2B; full distributions in Supplementary Table 2 and per-category tests in Supplementary Table 3). The shift was not explained by general differences in language production or acoustic properties of speech, which were comparable across questions and groups (Supplementary Table 1).

**Figure 2.**
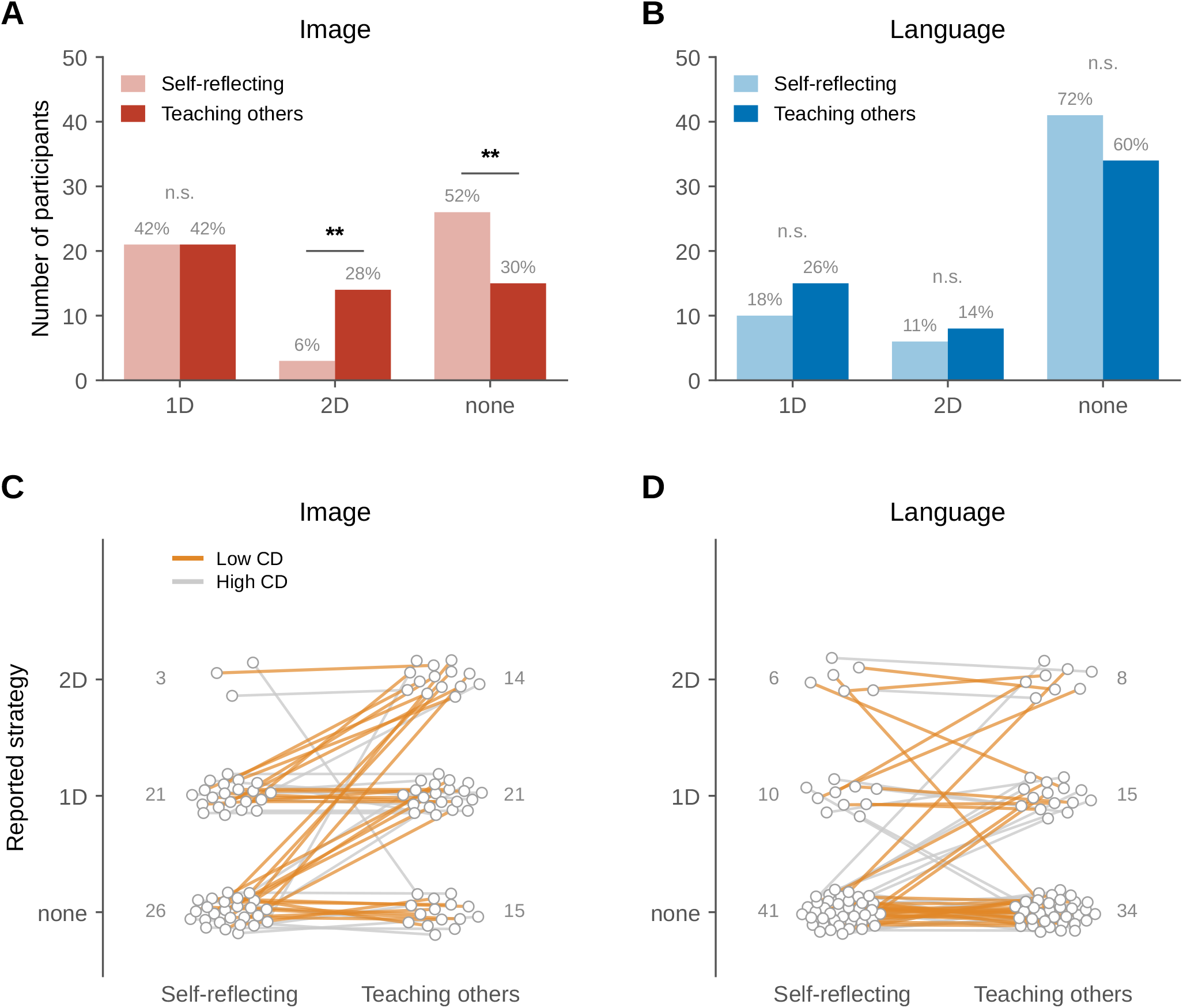
Teaching others externalizes spatial structure, selectively after image-based learning. A-B.Number of participants reporting a sequential (1D), spatial (2D), or no strategy when reflecting on their own memory (light bars) and when teaching others (solid bars), in the image-(A) and language-based (B) conditions. Percentages of the group are given above the bars. **C-D**. Individual transitions in reported strategy from self-reflection to teaching others in the image-(C) and language-based (D) conditions. Each participant is represented by one dot per column, placed according to the reported strategy; the two dots of the same participant are connected by a line coloured by cognitive disorganization (CD; O-LIFE) group defined by median split (low CD, orange; high CD, grey). Numbers next to each cluster give the number of participants in that category; the corresponding statistics are reported in the text. In (A) and (B), asterisks denote within-participant differences between self-reflection and teaching others (exact McNemar test; *\*p <* .05, *\*\*p <* .01); n.s., not significant.

### Cognitive disorganization and spatial descriptions

We next asked whether individual differences in schizotypy modulate this externalization. In a logistic regression predicting 2D strategy reports from group, question, age, gender, and all four O-LIFE subscales, CD was the only dimension associated with a lower likelihood of describing 2D structure (*β* = *−*0.544, *p* = .041; Supplementary Table 4, Supplementary Fig. 1); this association held with participant-clustered standard errors (*p* = .045) and attenuated in a random-intercept model accounting for the two reports per participant (*b* = *−*0.97, *p* = .070; Supplementary Table 5), and no other subscale showed a comparable negative effect, indicating specificity to CD rather than to schizotypy in general.

Because each participant provided both reports, we examined the same data as individual transitions between the two questions (Figure 2C–D). After image-based learning, participants with low CD (median split; *n* = 30) showed a marked increase in 2D descriptions from self-reflection to teaching (1/30 vs. 11/30; ten participants gained and none lost a 2D strategy; exact McNemar test, *p* = .002; Figure 2C), most of them moving from a sequential or unstructured account to a spatial one (verbatim examples in Supplementary Table 6). Participants with higher CD (*n* = 20) largely retained their initial category (2/20 vs. 3/20; exact McNemar *p* = 1.000). After language-based learning, transitions were infrequent and showed no systematic direction in either CD group (low CD: *n* = 32, 4/32 vs. 5/32, exact McNemar *p* = 1.000; high CD: *n* = 25, 2/25 vs. 3/25, exact McNemar *p* = 1.000; Figure 2D). Results survived Bonferroni correction (*α* = .0125). We emphasise, however, that these are within-group tests: the direct comparison between low- and high-CD participants in the image condition was not significant (11/30 vs. 3/20 at teaching, Fisher’s exact *p* = .118), and a logistic model with a continuous group *×* CD interaction likewise showed no reliable interaction (Supplementary Table 5); the model-implied probability of a 2D report as a continuous function of CD is shown in Supplementary Fig. 2. The categorical dissociation is therefore suggestive rather than conclusive, and we treat the continuous analysis reported next as the test of whether CD acts differently on the two learning modalities.

### Disorganization is expressed as tangential speech

Strategy coding captures what participants said about structure, but not how well their speech answered what they had been asked. We therefore computed a second, rater-free measure on the same recordings: the cosine similarity between the embedded question and each embedded sentence of the answer (semantic relevance; Figure 3A–B; Methods). Low values mean that a participant’s sentences drift away from the question they were asked, that is, that the answer is tangential to the question. Semantic relevance decreased from self-reflection to teaching in both conditions (image: *d*_*z*_ = *−*0.54; language: *d*_*z*_ = *−*0.63; both *p <* .001), with no difference between conditions (interaction *p* = .74), as expected when the task shifts from reporting a strategy to elaborating it for a listener (Supplementary Fig. 3). Critically, semantic relevance was related to CD in a modality-specific way. Following image-based learning, participants with higher CD gave answers that tracked the question less closely (*r* = *−*0.47, *p <* .001), whereas no such relation was observed following language-based learning (*r* = 0.20, *p* = .14; group *×* CD interaction, *p <* .001; Figure 3C– D). This interaction was robust to controlling for the other O-LIFE subscales, age, gender, and the language of the report, and held in a sentence-level mixed model with participants as random effects (image: *b* = *−*0.63 per SD of CD, *p <* .001; language: *b* = 0.26, *p* = .054; interaction *p <* .001).

**Figure 3.**
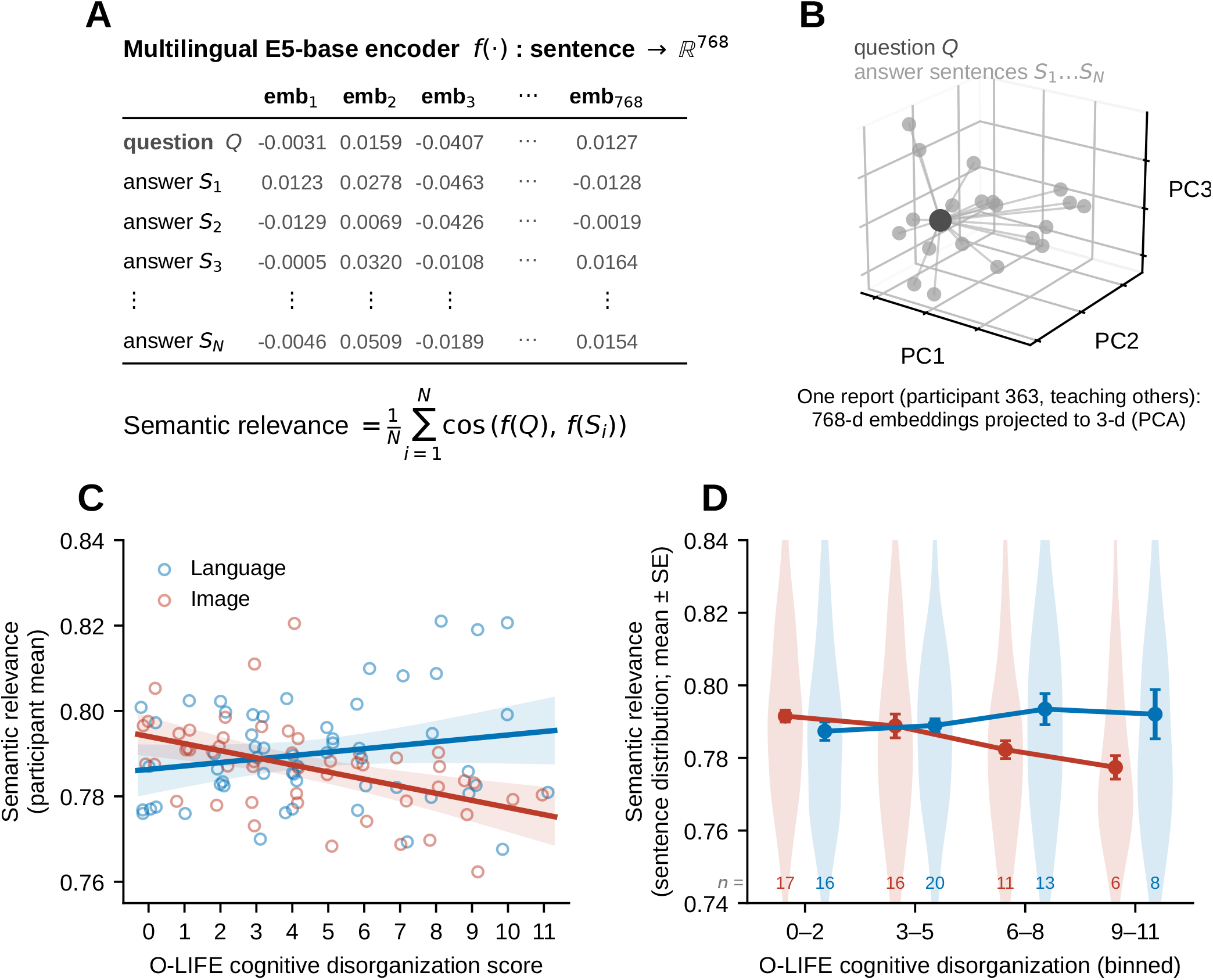
After image-based learning, cognitive disorganization is associated with answers that stray from the question asked. A.Each sentence *S*_*i*_ of a report and the question *Q* were encoded with a multilingual sentence encoder (multilingual-E5-base, 768 dimensions); semantic relevance is the mean cosine similarity between the question and the sentences of the answer, so that low values indicate an answer that is tangential to the question. **B**. Example report (participant 363, teaching others): the question (dark) and the 20 sentences of the report (grey) projected from 768 to 3 dimensions by principal component analysis; lines connect each sentence to the question. **C**. Semantic relevance (participant mean across both reports; one circle per participant) as a function of O-LIFE cognitive disorganization (CD) in the image-(red) and language-based (blue) conditions, with linear fits and 95% confidence bands. **D**. Same data with CD binned; violins show the distribution of sentence-level relevance, points show mean *±* SE of participant means, and numbers give participants per bin. Mixed-model slopes are from a sentence-level model with random intercepts for participants.

Two features of this effect distinguish it from the strategy result and clarify what it measures. First, it was present in both reports taken separately (self-reflection *r* = *−*0.33, *p* = .021; teaching *r* = *−*0.46, *p <* .001; Supplementary Fig. 4), and CD was unrelated to the change between them (*r* = *−*0.05, *p* = .76). The tangentiality associated with CD is therefore a property of how these participants answered questions after visual learning, not a failure of the teaching manipulation to take effect. Second, it is not a by-product of speaking more or less: CD was essentially unrelated to word count, sentence count, speech rate, duration and pauses, and the interaction was unchanged when these were added as covariates (Supplementary Table 5). The measure is estimated on a continuous scale without a median split, is robust to the language of the report, and is visible at the level of individual sentences as well as whole reports (Supplementary Fig. 4).

## Discussion

Explaining learned material to another person brought two-dimensional structure into natural speech, and did so selectively after visual learning; on the same recordings, higher cognitive disorganization was associated with answers that stayed further from the question asked, again selectively after visual learning. That explanation, but not self-reflection, elicits spatial descriptions suggests that relational structure is present in memory but not routinely verbalized: it surfaces when speech must serve a listener. That it surfaces only after image-based learning is consistent with the idea that the two learning formats yield representations of different depth. When relations are stated in sentences, participants can recall the sentences without ever composing them into a space; when relations must be inferred from images, the resulting representation is spatial from the outset and is available to be described.

The two readouts therefore describe different things that point the same way. Explanation brings two-dimensional structure into speech, and only after visual learning (Figure 2); after visual learning, higher CD is accompanied by answers that stay further from the question (Figure 3). Note that the two directions of change are not in conflict: relevance falls for everyone when the task turns to teaching, which reflects elaboration rather than disorganization, whereas the CD effect is a between-person difference that is present in both questions and does not act on that change.

Taken together, these results place the shallow cognitive map account on a new kind of evidence. The hypothesis has so far been supported by task performance and neural measures^1,2^; here the same signature appears in what people say, in ordinary language produced for a listener. Two features of the pattern constrain its interpretation. First, CD was associated with a lower probability of describing 2D structure overall (Supplementary Table 4, Supplementary Fig. 1) and with answers that stayed further from the question asked, rather than with a general reduction in speaking or in structure of any kind: it did not shift reports along the ordered scale from no strategy through sequential to spatial descriptions (Supplementary Table 5), and it was unrelated to how much participants said (Supplementary Table 5). What appears to be compromised is the two-dimensional organization itself rather than verbal output in general. Second, the effect was confined to the image-based condition. Because both conditions conveyed identical relations and produced comparable final accuracy (Supplementary Table 1), the difference is unlikely to reflect how well the material was learned; it points instead to the stage at which relational structure must be internally generated rather than supplied by language, consistent with our earlier finding in task performance ^13^.

That spatial description also tracked independent behavioral measures supports this reading. Participants who described spatial strategies during self-reflection showed better map-location memory of the conceptual space and higher working memory accuracy (Supplementary Fig. 6), indicating that spatial language reflects differences in conceptual organization rather than in verbal style. In this respect the present findings connect the literature on multidimensional relational codes for abstract knowledge^6–9,14^ to work showing that language production carries information about disruptions in underlying semantic structure^10,11^.

The practical implications are worth stating plainly. The teaching prompt is a three-minute, language-only probe: it requires no psychometric task, no reading, no motor response, and no equipment beyond a microphone, and the semantic-relevance measure is computed automatically, without raters, in whatever language the speaker prefers. This profile is suited to settings where structured cognitive testing is difficult, including remote and digital assessment, and to dimensional approaches that track subclinical variation over time. Because the measure indexes a specific construct, the organization of relational knowledge, rather than global impairment, it may complement existing speech markers of psychosis risk, which have largely targeted coherence and syntactic complexity^10,11^. Establishing clinical utility will require testing whether the same readout tracks symptomatic disorganization in patients, and whether it changes with treatment.

## Limitations

Several limitations qualify these conclusions. The sample was non-clinical and varied in schizotypy rather than diagnosis, so whether the same signature appears in psychosis remains to be tested. The design was cross-sectional and used a single two-dimensional structure; whether the effect generalizes to higher-dimensional or non-orthogonal spaces is unknown. The categorical strategy analyses relied on a median split for visualization, and the modality-specific effect of CD on strategy reports was not significant when tested continuously; the dissociation rests on the continuous semantic-relevance analysis, and a replication with a larger sample would be needed to establish it for the categorical readout as well. Finally, reports were produced in several languages. Language of report was included as a covariate throughout, and the results were unchanged when all reports were translated into English and re-embedded (Supplementary Fig. 5), which makes a language artefact unlikely; a sample tested in a single language would nevertheless be a cleaner design. A direct test of the account would combine this paradigm with a clinical sample and with neural measures of map structure, asking whether the speech readout and the neural readout are disrupted together.

## Methods

### Participants

Participants were right-handed adults (*≥*18 years) recruited via social media from the university community in Zurich, most of whom were university students. Inclusion required normal or corrected-to-normal vision, English proficiency at C1 level or above, and no prior education or training in psychology. A total of 111 participants completed the study, which lasted 2–3 hours (mean 2 h 10 min), and each received 40 CHF as compensation. Participants ranged in age from 18 to 64 years. Audio recordings were unavailable for four participants because of equipment failure, so all analyses of verbal reports were conducted on the remaining 107 participants (50 assigned to the image-based and 57 to the language-based condition). Sample characteristics, including age, gender, years of education, English proficiency, and language of the verbal report, are reported by group in Supplementary Table 1. Written informed consent was obtained from all participants before the experiment. The study adhered to the principles of the Declaration of Helsinki and was approved by the Cantonal Ethics Committee Zurich (Kantonale Ethikkommission Zürich; Protocol No. 2024-01314).

The learning and testing tasks of this dataset, but not the verbal reports, are reported in a separate paper^13^; the audio recordings analyzed here were explicitly not included in that report.

Self-reported gender is reported in Supplementary Table 1 and was included as a covariate in the regression models; the study was not designed to test for effects of sex or gender, and no such analyses were preregistered.

### Conceptual structure and learning phase

Participants learned six food items, each paired with a unique combination of two orthogonal conceptual dimensions: the age group of the person eating the food and the time of day at which it was eaten. Each dimension comprised seven graded levels (age group: toddler, child, adolescent, adult, middle-aged, elderly, old; time of day: dawn, morning, noon, afternoon, nightfall, evening, night), so that the two dimensions together define a two-dimensional conceptual space (Figure 1A). Participants were never shown the space itself, and terms such as “map” or “two-dimensional space” were never used. Participants were randomly assigned to an image-based or a language-based condition, which conveyed identical conceptual relationships either through cartoon faces and symbolic icons or through English words and short sentences (Figure 1B).

The learning phase comprised three tasks, described in full in our previous report using the same dataset^13^. In the *ordering task*, participants judged the relative order of two randomly selected levels of the same dimension (for example, whether “toddler” is younger than “adolescent”); training continued until participants reached at least 85% accuracy over the most recent 20 trials, after which they arranged all seven levels of the dimension from smallest to largest. The order in which the two dimensions were learned was randomized across participants. In the *association learning task*, a target age–time combination was displayed with six candidate food items below it; participants selected the correct item with the mouse and received immediate feedback, and training continued until they reached at least 85% accuracy over the most recent 20 trials. In the *navigation task*, a food item was displayed and participants selected the corresponding age group and time of day from the full set of graded levels; feedback was immediate, a trial counted as correct only if both selections were correct on the first attempt, and training continued until participants reached 12 consecutive correct trials. Both groups reached comparable accuracy in all three tasks, although the language-based group required more trials and more time to reach criterion (Supplementary Table 1).

After the learning phase and before the verbal reports, participants completed testing-phase tasks (similarity ratings, a reward-based choice task, and a placement task) and a post-experiment assessment; those data are reported elsewhere^13^ and are not analyzed here, except for the placement task, the n-back task, and the mental rotation task, which provide the cognitive measures reported in Supplementary Table 1.

### Verbal report task

After the learning phase, participants completed a verbal report task programmed in PsychoPy (v2021.2.3), with audio captured through the computer’s input device using the sounddevice and wavio libraries at a sampling rate of 48 kHz and 16-bit depth. Participants were told that they would answer two questions verbally, that they would have three minutes for each, and that they could speak in any language in which they felt comfortable, with their mother tongue recommended. They were asked to think about and prepare each answer before starting, to press a key when ready, and to try to speak for the full three minutes; a countdown timer was displayed throughout. Recording lasted 180 s per question irrespective of when the participant stopped speaking.

The two questions were presented in fixed order. Self-reflection: “Think back to how you remembered the six combinations. That is, how age group and time were connected to food. Did you use any memory strategies? Did you find it difficult to remember them?” Teaching others: “Imagine you are talking to someone who has never learned these combinations before. What would you say to help them better remember the six combinations?”

### Questionnaires and cognitive tasks

Before the experiment, participants provided demographic information and completed the short form of the Oxford–Liverpool Inventory of Feelings and Experiences (sO-LIFE) ^15^, which yields four subscales (Unusual Experiences, Cognitive Disorganization, Introvertive Anhedonia, and Impulsive Nonconformity), the Big Five Inventory ^16^, and the Positive and Negative Affect Schedule^17^. Participants also reported their hunger level and the time since their last meal, to control for appetite-related confounds arising from the food stimuli. After the experiment, participants completed an n-back task^18^ indexing working memory and a mental rotation task^19^ indexing spatial transformation ability, each lasting approximately 7–8 minutes. The n-back measure reported here is corrected accuracy (hits minus false alarms) and the mental rotation measure is the number of correct responses. Descriptive statistics for all measures are given in Supplementary Table 1.

### Transcription and strategy classification

Audio recordings were transcribed with Whisper (large-v3) in the language spoken by the participant. Two independent raters classified each report into one of three categories: 2D, if the participant described a two-dimensional spatial structure with relationships along two distinct dimensions, such as a grid or plane; 1D, if the participant organized items along a single dimension, such as a timeline or a linear sequence; and none, if no explicit organizational strategy was described. Of the 214 reports, 19 showed initial disagreement between raters (Cohen’s *κ* = 0.85). These were reviewed independently a second time and resolved by consensus discussion, and the final consensus classification was used in all analyses.

### Semantic relevance

Transcripts were split into sentences with the spaCy (v3.8.11) multilingual sentencizer, yielding 3,380 sentences (4–34 per report). Each sentence and each question prompt was encoded with the multilingual sentence encoder multilingual-E5-base (768 dimensions, L2-normalized), using the “query:” prefix for the question and the “passage:” prefix for the sentences, as recommended for this model. Semantic relevance of a report was defined as the mean cosine similarity between the embedded question and the embedded sentences of the answer to it; low values indicate an answer that drifts away from the question asked. Because the encoder is multilingual, reports were analyzed in the language in which they were produced, and the language of the report was included as a covariate in all regression models. As a robustness analysis, all 214 reports were machine-translated into English (Google Translate via the deep-translator Python package) and the translations were checked manually; the translated reports were split into sentences and embedded with the same encoder and prefixes, and all results were unchanged (Supplementary Fig. 5).

### Statistical analysis

All statistical tests were two-sided unless stated otherwise. Within-participant changes in reported strategy between the two questions were tested with exact McNemar tests on the discordant pairs; between-group comparisons used Fisher’s exact tests. Logistic regression models predicted the probability of reporting a 2D strategy from learning condition, question, age, gender, and all four sO-LIFE subscales; reference levels were the image condition, self-reflection, and female. For the analyses shown in Figure 2C–D, participants were divided into low- and high-CD groups by a median split of the Cognitive Disorganization subscale (low CD defined as a score at or below the median). Semantic relevance was analyzed both at the level of participant means, with Pearson correlations and linear models including the other sO-LIFE subscales, age, gender, and language of report as covariates, and at the level of individual sentences, with linear mixed models including random intercepts for participants. Where multiple comparisons were conducted, Bonferroni correction was applied (*α* = .0125). Statistical details for each test, including the test used, the value of *n*, and what *n* represents, are reported in the corresponding figure legends, in the Results text, and in Supplementary Tables 1–4. Analyses were conducted in MATLAB R2023b and in Python 3.11.5.

### Preregistration

The parent study was preregistered on AsPredicted (#z8tn-kjvw; https://aspredicted.org/z8tn-kjvw.pdf). The preregistration specified the learning paradigm, the two learning conditions, the schizotypy and cognitive measures, and a target sample of at least 80 participants (40 per group) based on an a priori power analysis in G*Power. The verbal report task and all analyses of speech reported here were not part of the preregistration and are therefore exploratory.

## Data availability

De-identified transcripts (original language and English translations), strategy ratings, sentence-level semantic relevance values, and questionnaire scores are available athttps://github.com/xiaoyanwu2024/verbalLanguage-ssd. Raw audio recordings are not shared because they are potentially identifying; they are available from the corresponding authors on request under a data use agreement.

## Code availability

All code for the analyses and figures reported in this paper is available at https://github.com/xiaoyanwu2024/verbalLanguage-ssd.

## Acknowledgements

This work was funded by the European Research Council (ERC), EU Horizon Europe programme (grant agreement no. 101118756), as part of the ERC Synergy project DELTA-LANG. X.W. was supported by a UZH Postdoc Grant from the University of Zurich (grant no. FK-26-058). We thank Ray Dolan for insightful discussions.

## Author contributions

Conceptualization, P.H. and X.W.; methodology, software, validation, formal analysis, investigation, and resources, X.W.; data curation, X.W. and P.W.; writing–original draft, X.W.; writing–review & editing, X.W., P.W., W.O., W.H., I.E.S., and P.H.; visualization, X.W.; supervision, P.H.; project administration, P.H. and X.W.; funding acquisition, P.H. and X.W.

## References

1. Musa, A., Khan, S., Mujahid, M., and El-Gaby, M. (2022). The shallow cognitive map hypothesis: A hippocampal framework for thought disorder in schizophrenia. Schizophrenia 8, 34. doi: 10.1038/s41537-022-00247-7.

2. Nour, M.M., Liu, Y., El-Gaby, M., McCutcheon, R.A., and Dolan, R.J. (2025). Cognitive maps and schizophrenia. Trends in Cognitive Sciences 29, 184–200. doi: 10.1016/j.tics.2024.09.011.

3. Tolman, E.C. (1948). Cognitive maps in rats and men. Psychological review 55, 189. doi: 10.1037/h0061626.

4. Bellmund, J.L., Gärdenfors, P., Moser, E.I., and Doeller, C.F. (2018). Navigating cognition: Spatial codes for human thinking. Science 362, eaat6766. doi: 10.1126/science.aat6766.

5. Behrens, T.E., Muller, T.H., Whittington, J.C., Mark, S., Baram, A.B., Stachenfeld, K.L., and Kurth-Nelson, Z. (2018). What is a cognitive map? Organizing knowledge for flexible behavior. Neuron 100, 490–509. doi: 10.1016/j.neuron.2018.10.002.

6. Constantinescu, A.O., O’Reilly, J.X., and Behrens, T.E. (2016). Organizing conceptual knowledge in humans with a gridlike code. Science 352, 1464–1468. doi: 10.1126/science.aaf0941.

7. Theves, S., Fernandez, G., and Doeller, C.F. (2019). The hippocampus encodes distances in multidimensional feature space. Current Biology 29, 1226–1231. doi: 10.1016/j.cub.2019.02.035.

8. Viganó, S., and Piazza, M. (2020). Distance and direction codes underlie navigation of a novel semantic space in the human brain. Journal of Neuroscience 40, 2727–2736. doi: 10.1523/JNEUROSCI.1849-19.2020.

9. Zheng, X.Y., Hebart, M.N., Grill, F., Dolan, R.J., Doeller, C.F., Cools, R., and Garvert, M.M. (2024). Parallel cognitive maps for multiple knowledge structures in the hippocampal formation. Cerebral Cortex 34, bhad485. doi: 10.1093/cercor/bhad485.

10. Nour, M.M., McNamee, D.C., Liu, Y., and Dolan, R.J. (2023). Trajectories through semantic spaces in schizophrenia and the relationship to ripple bursts. Proceedings of the National Academy of Sciences 120, e2305290120. doi: 10.1073/pnas.2305290120.

11. Fradkin, I., Adams, R.A., Siegelman, N., Moran, R., and Dolan, R.J. (2024). Latent mechanisms of language disorganization relate to specific dimensions of psychopathology. Nature Mental Health 2, 1486–1497. doi: 10.1038/s44220-024-00351-w.

12. Lombrozo, T. (2006). The structure and function of explanations. Trends in cognitive sciences 10, 464–470. doi: 10.1016/j.tics.2006.08.004.

13. Wu, X., Rabe, F., Wu, P., Edkins, V., Pauli, Y., Omlor, W., Misra, A.R., Theves, S., Hinzen, W., Sommer, I.E. et al. (2026). Disorganization impairs cognitive maps built from visual inputs. Schizophrenia Bulletin. In press.

14. Garvert, M.M., Saanum, T., Schulz, E., Schuck, N.W., and Doeller, C.F. (2023). Hippocampal spatio-predictive cognitive maps adaptively guide reward generalization. Nature Neuroscience 26, 615–626. doi: 10.1038/s41593-023-01283-x.

15. Mason, O., and Claridge, G. (2006). The Oxford-Liverpool inventory of feelings and experiences (O-LIFE): Further description and extended norms. Schizophrenia Research 82, 203–211. doi: 10.1016/j.schres.2005.12.845.

16. McCrae, R.R., and Costa, J., Paul T. (1999). A five-factor theory of personality. In L.A. Pervin, and O.P. John, eds. Handbook of Personality: Theory and Research pp. 139–153. New York: Guilford Press.

17. Watson, D., Clark, L.A., and Tellegen, A. (1988). Development and validation of brief measures of positive and negative affect: The PANAS scales. Journal of Personality and Social Psychology 54, 1063–1070. doi: 10.1037/0022-3514.54.6.1063.

18. Colwell, M., Tagomori, H., Martens, M., Murphy, S., and Harmer, C. (2023). N-back (Oxford) — fMRI and non-scanner release. Zenodo. doi: 10.5281/zenodo.8003407.

19. Shepard, R.N., and Metzler, J. (1971). Mental rotation of three-dimensional objects. Science 171, 701–703. doi: 10.1126/science.171.3972.701.

