## Supporting Information for "Teaching others reveals hidden structure in conceptual cognitive maps"

#### Supplementary figures

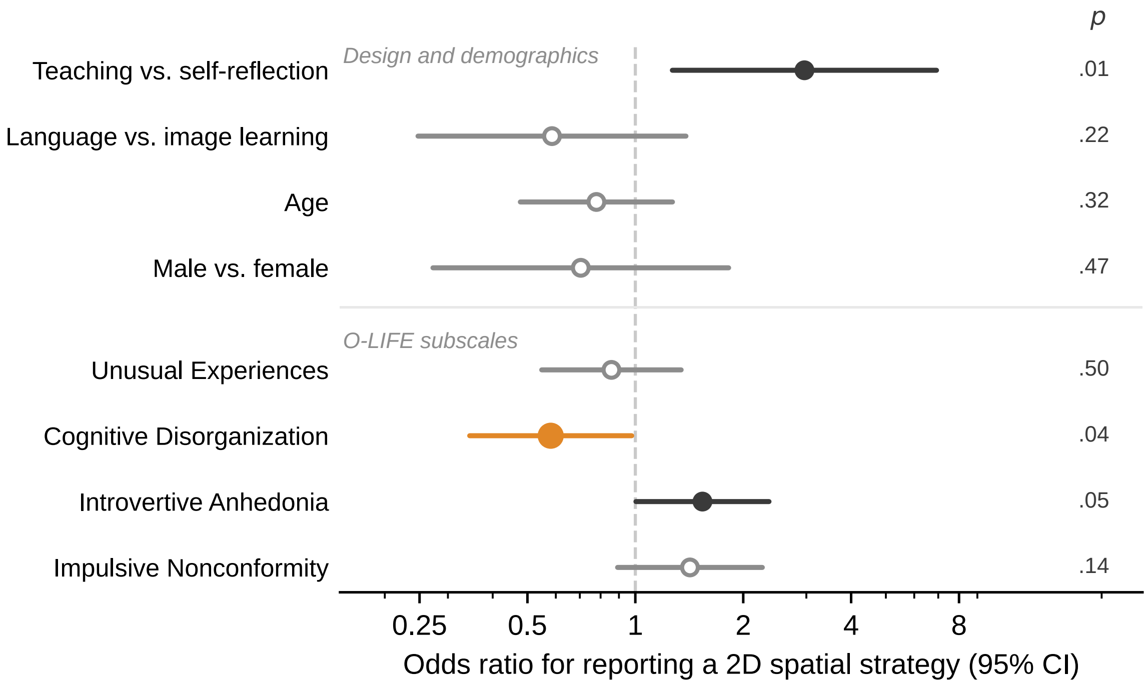


Supplementary Fig. 1 | Logistic regression predicting a 2D spatial strategy report. Odds ratios with 95% confidence intervals from the model in Supplementary Table 4 (214 reports from 107 participants). Filled markers indicate $p < .05$; the O-LIFE subscales are $z$-scored, so their odds ratios refer to a one-SD change. Cognitive Disorganization (orange) is the only subscale associated with a lower probability of reporting 2D structure; Introvertive Anhedonia shows an association in the opposite direction. Confidence intervals are from the unadjusted model; repeated-measures corrections are reported in Supplementary Table 5.

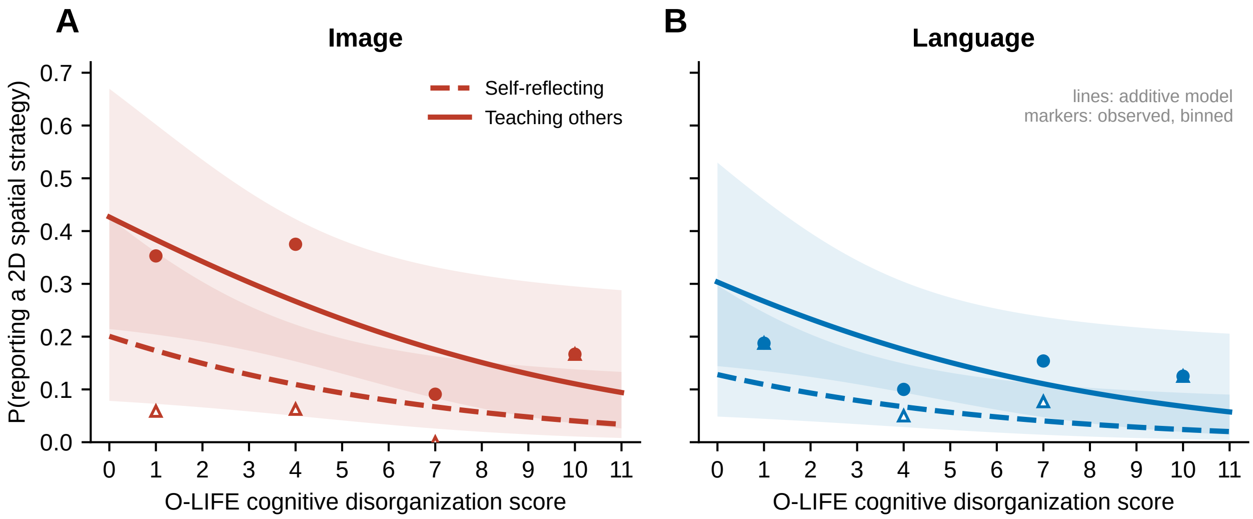


Supplementary Fig. 2 | Probability of reporting a 2D spatial strategy as a continuous function of cognitive disorganization. Lines give the probability implied by the logistic model of Supplementary Table 4 with all other covariates at their sample means and gender at the reference level, for self-reflection (dashed) and teaching others (solid); shaded bands are 95% confidence intervals. Markers give observed proportions in bins of the CD score (0–2, 3–5, 6–8, 9–11) for bins containing at least five reports. **A.** Image-based learning. **B.** Language-based learning. The model is additive in CD; a version including a group × CD interaction is reported in Supplementary Table 5 and did not improve on it.

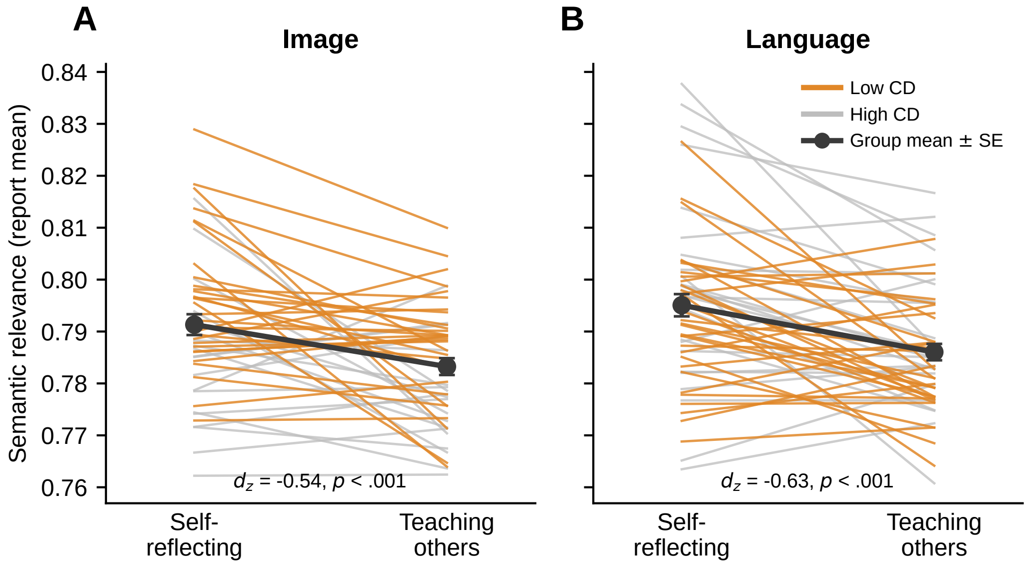


Supplementary Fig. 3 | Semantic relevance of each participant's two reports. Each line joins one participant's self-reflection and teaching report, coloured by cognitive disorganization group (median split). Black markers give the group mean ± SE. **A.** Image-based learning. **B.** Language-based learning. Relevance decreased from self-reflection to teaching in both conditions, consistent with explanatory speech moving beyond the literal material; the two conditions did not differ in the size of this decrease.

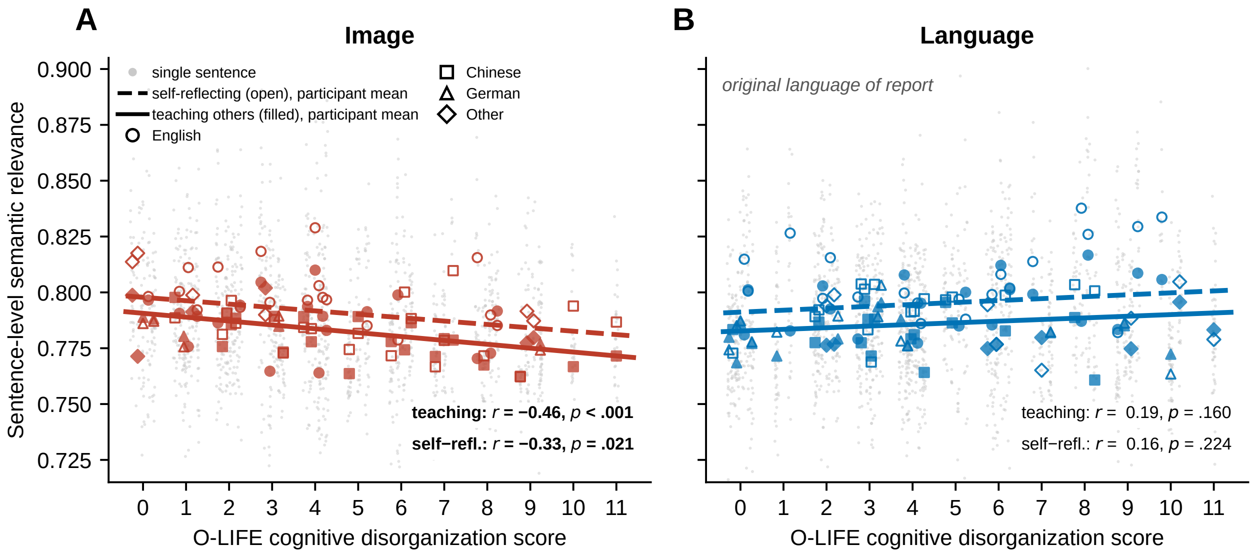


Supplementary Fig. 4 | Sentence-level semantic relevance as a function of cognitive disorganization, by question. Each small grey dot is one sentence, arranged in a narrow column per participant. Markers give participant means for the self-reflection (open) and teaching (filled) reports, with marker shape indicating the language of the report; dashed and solid lines are linear fits to the participant means of the two questions, with Pearson correlations given in each panel. **A.** Image-based learning: the negative relation with CD is present for both questions. **B.** Language-based learning: no relation for either question.

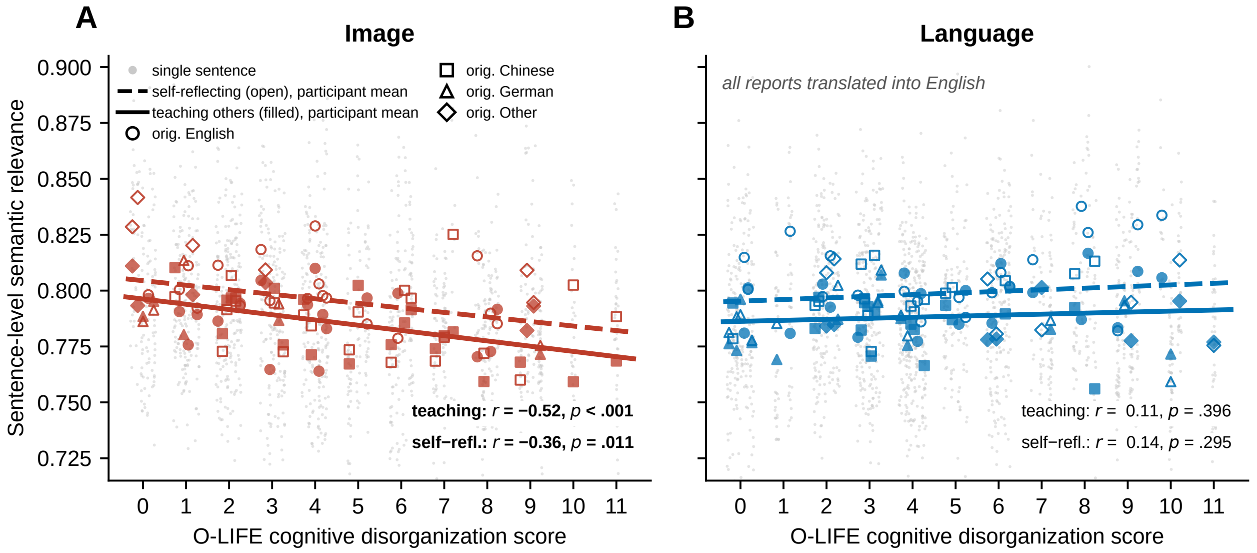


Supplementary Fig. 5 | The analysis of Supplementary Fig. 4 repeated on English translations of all reports. All 214 reports were translated into English, split into sentences, and re-embedded with the same encoder, removing any residual influence of the language of the report on the embeddings; marker shapes indicate the original language of each report, as in Supplementary Fig. 4. The pattern is unchanged (image condition: self-reflection $r = -0.36$, $p = .011$; teaching $r = -0.52$, $p < .001$; language condition: both $p > .17$; group × CD interaction $b = 0.0085$, $p = 2.6 \times 10^{-4}$), and participant-level relevance computed from the original-language and translated texts correlated at $r = .92$. The effect in Figure 3 is therefore not attributable to the multilingual embedding treating languages differently.

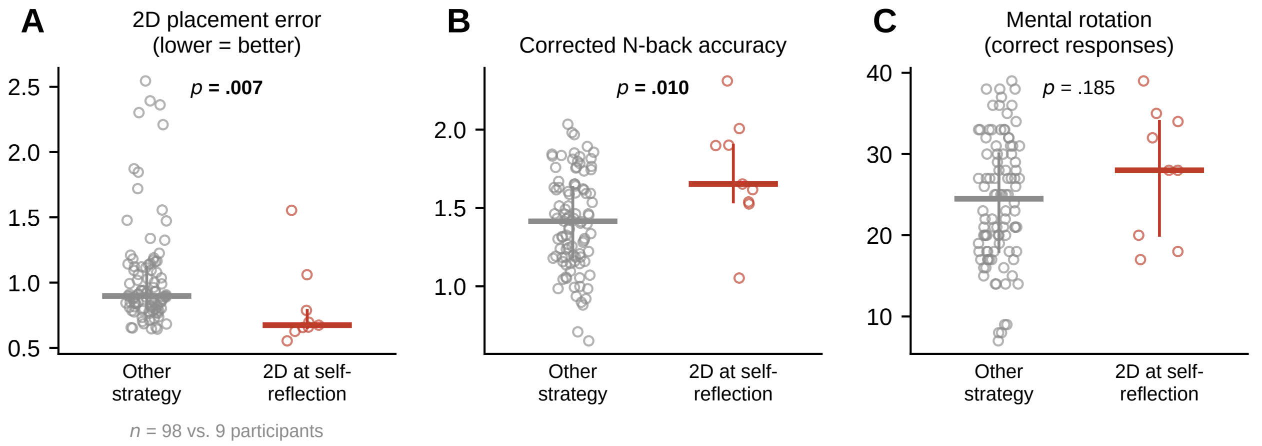


Supplementary Fig. 6 | Cognitive performance of participants who did and did not describe a 2D spatial strategy during self-reflection. Open circles are individual participants (98 who described another strategy, 9 who described 2D structure); horizontal bars give the median and vertical bars the interquartile range. **A.** 2D placement error, the mean Euclidean distance between dragged and true item locations. **B.** Corrected N-back accuracy. **C.** Mental rotation performance. $p$ values are from two-sided Wilcoxon rank-sum tests, uncorrected. Note the small number of participants who spontaneously described 2D structure during self-reflection; these comparisons are exploratory.

#### Supplementary tables

**Supplementary Table 1 | Participant characteristics, task performance, and speech-production features in the image- and language-based learning groups.**

| | Image ($n$ = 50) | Language ($n$ = 57) | Image vs. Language |
|---|---|---|---|
| *Demographics* | | | |
| Age, years | 26.61 (4.70) | 27.51 (6.85) | $t = -0.78$, $p$ = .436, $d = -0.15$ |
| Gender, female/male | 33/17 | 41/16 | $\chi^2$ = 0.44, $p$ = .508, $V$ = 0.06 |
| Education, years | 16.46 (1.88) | 16.60 (2.47) | $t = -0.32$, $p$ = .751, $d = -0.06$ |
| English proficiency, native-like/high (C1)/moderate (B1–B2) | 10/38/2 | 11/44/2 | $\chi^2$ = 0.03, $p$ = .986, $V$ = 0.02 |
| Language of verbal report, English/Chinese/German/other | 18/21/5/6 | 20/16/13/8 | $\chi^2$ = 4.18, $p$ = .242, $V$ = 0.20 |
| *Schizotypy (O-LIFE)* | | | |
| Unusual Experiences | 3.14 (2.49) | 3.46 (2.87) | $t = -0.60$, $p$ = .547, $d = -0.12$ |
| Cognitive Disorganization | 4.26 (3.05) | 4.53 (3.07) | $t = -0.45$, $p$ = .654, $d = -0.09$ |
| Introvertive Anhedonia | 2.68 (1.99) | 2.61 (2.09) | $t$ = 0.17, $p$ = .868, $d$ = 0.03 |
| Impulsive Nonconformity | 1.84 (1.35) | 2.79 (1.87) | $t = -2.98$, $p$ = .004, $d = -0.58$ |
| Total score | 11.92 (6.19) | 13.39 (6.73) | $t = -1.17$, $p$ = .246, $d = -0.23$ |
| *Personality and affect* | | | |
| Extraversion (BFI) | 25.40 (4.98) | 25.84 (5.59) | $t = -0.43$, $p$ = .668, $d = -0.08$ |
| Agreeableness (BFI) | 31.26 (3.67) | 30.47 (4.03) | $t$ = 1.05, $p$ = .297, $d$ = 0.20 |
| Conscientiousness (BFI) | 29.84 (5.64) | 29.37 (5.52) | $t$ = 0.44, $p$ = .664, $d$ = 0.08 |
| Neuroticism (BFI) | 23.22 (5.60) | 23.65 (6.59) | $t = -0.36$, $p$ = .719, $d = -0.07$ |
| Openness (BFI) | 36.06 (6.04) | 38.23 (5.59) | $t = -1.93$, $p$ = .056, $d = -0.37$ |
| Positive affect (PANAS) | 31.96 (5.84) | 31.40 (6.53) | $t$ = 0.46, $p$ = .645, $d$ = 0.09 |
| Negative affect (PANAS) | 22.04 (7.00) | 20.70 (7.06) | $t$ = 0.98, $p$ = .328, $d$ = 0.19 |
| *Learning-phase performance* | | | |
| Ordering task, accuracy | 0.982 (0.029) | 0.971 (0.037) | $t$ = 1.73, $p$ = .087, $d$ = 0.33 |
| Ordering task, total response time, s | 69.9 (25.0) | 91.8 (33.2) | $t = -3.82$, $p <$ .001, $d = -0.74$ |
| Association learning, accuracy | 0.927 (0.100) | 0.895 (0.129) | $t$ = 1.40, $p$ = .164, $d$ = 0.27 |
| Association learning, number of trials | 41.2 (23.7) | 75.2 (80.2) | $t = -2.89$, $p$ = .005, $d = -0.56$ |
| Association learning, total response time, s | 128.7 (68.7) | 237.0 (201.9) | $t = -3.61$, $p <$ .001, $d = -0.70$ |
| Navigation task, accuracy | 0.958 (0.026) | 0.965 (0.053) | $t = -0.92$, $p$ = .361, $d = -0.18$ |
| Navigation task, total response time, s | 177.3 (140.8) | 186.1 (174.2) | $t = -0.28$, $p$ = .778, $d = -0.05$ |
| *Cognitive performance* | | | |
| Mental rotation, correct responses | 23.5 (7.5) | 25.3 (7.8) | $t = -1.24$, $p$ = .216, $d = -0.24$ |
| Corrected N-back accuracy | 1.40 (0.33) | 1.46 (0.30) | $t = -1.01$, $p$ = .312, $d = -0.20$ |
| *Speech production*[a] | | | |
| Word count | | | |
| Self-reflection | 216.3 (160.1) | 230.8 (136.5) | $t = -0.51$, $p$ = .614, $d = -0.10$ |
| Teaching others | 211.8 (162.0) | 229.2 (139.4) | $t = -0.60$, $p$ = .552, $d = -0.12$ |
| $\Delta$ (Teaching − Self-reflection) | $-4.50$, $p$ = .439 | $-1.65$, $p$ = .829 | $t = -0.29$, $p$ = .771 |
| Sentence count | | | |
| Self-reflection | 17.1 (10.3) | 16.8 (6.3) | $t$ = 0.20, $p$ = .839, $d$ = 0.04 |
| Teaching others | 15.5 (6.4) | 16.9 (7.3) | $t = -1.05$, $p$ = .298, $d = -0.20$ |
| $\Delta$ (Teaching − Self-reflection) | $-1.68$, $p$ = .184 | 0.05, $p$ = .947 | $t = -1.20$, $p$ = .232 |
| Speech duration, s | | | |
| Self-reflection | 107.6 (25.9) | 94.8 (31.1) | $t$ = 2.30, $p$ = .024, $d$ = 0.44 |
| Teaching others | 103.3 (28.5) | 92.7 (29.7) | $t$ = 1.88, $p$ = .063, $d$ = 0.36 |
| $\Delta$ (Teaching − Self-reflection) | $-4.29$, $p$ = .153 | $-2.12$, $p$ = .373 | $t = -0.58$, $p$ = .565 |
| Pause count | | | |
| Self-reflection | 70.3 (20.4) | 77.8 (29.4) | $t = -1.51$, $p$ = .133, $d = -0.29$ |
| Teaching others | 66.8 (20.8) | 74.6 (27.9) | $t = -1.62$, $p$ = .107, $d = -0.31$ |

*continued on next page*

*Supplementary Table 1, continued*

| | Image ($n$ = 50) | Language ($n$ = 57) | Image vs. Language |
|---|---|---|---|
| $\Delta$ (Teaching − Self-reflection) | −3.54, $p$ = .198 | −3.23, $p$ = .117 | $t$ = −0.09, $p$ = .926 |
| Speech rate | | | |
| Self-reflection | 3.93 (0.50) | 3.94 (0.49) | $t$ = −0.12, $p$ = .902, $d$ = −0.02 |
| Teaching others | 3.87 (0.62) | 3.92 (0.53) | $t$ = −0.46, $p$ = .644, $d$ = −0.09 |
| $\Delta$ (Teaching − Self-reflection) | −0.06, $p$ = .346 | −0.02, $p$ = .648 | $t$ = −0.52, $p$ = .605 |

*Note.* Values are mean (SD) unless otherwise indicated. Continuous measures were compared with independent-samples $t$ tests (effect size: Cohen's $d$), categorical measures with $\chi^2$ tests (effect size: Cramér's $V$). BFI = Big Five Inventory; PANAS = Positive and Negative Affect Schedule. Learning-phase measures are from the training session preceding the verbal reports. Impulsive Nonconformity differed between groups and was therefore included, together with the other O-LIFE subscales, as a covariate in all regression models. Accuracy in all learning-phase tasks was comparable between groups, but the language-based group required more trials and more time in the ordering and association-learning tasks. [a] For each speech feature, the first two rows give group means for the self-reflection and teaching-others reports. The row $\Delta$ gives the within-condition mean change (Teaching − Self-reflection) with the uncorrected $p$ value of a paired $t$ test; the right-hand column of that row tests whether this change differed between learning conditions (group × question interaction). No feature differed between self-reflection and teaching, and no feature showed a significant interaction. Speech duration was longer in the image-based group during self-reflection, but the change from self-reflection to teaching did not differ between groups. Semantic relevance of the reports to the learned material is analyzed separately in Figure 3 of the main text.

Supplementary Table 2 | Verbal strategy distribution by learning condition and question type. Values are $n$ (%).

| **Condition** | **Question** | **None** | **1D** | **2D** |
|---|---|---|---|---|
| Image ($n = 50$) | Self-reflection | 26 (52.0%) | 21 (42.0%) | 3 (6.0%) |
| | Teaching others | 15 (30.0%) | 21 (42.0%) | 14 (28.0%) |
| Language ($n = 57$) | Self-reflection | 41 (71.9%) | 10 (17.5%) | 6 (10.5%) |
| | Teaching others | 34 (59.6%) | 15 (26.3%) | 8 (14.0%) |

*Note.* Strategy categories: None = no identifiable spatial structure; 1D = one-dimensional (sequential) strategy; 2D = two-dimensional spatial strategy. Each row sums to 100%.

Supplementary Table 3 | Binary analysis of 2D strategy reports (exact McNemar test).

| Group | Self-reflection ($n$) | Teaching others ($n$) | Self-reflection (%) | Teaching others (%) | $p$ |
|---|---|---|---|---|---|
| Image | 3/50 | 14/50 | 6.0% | 28.0% | .003 |
| Language | 6/57 | 8/57 | 10.5% | 14.0% | .688 |

*Note.* $n$ values indicate the number of participants reporting a 2D spatial strategy out of the total in each condition. $p$ values are from exact McNemar tests for within-subject change from self-reflection to teaching others.

Supplementary Table 4 | Logistic regression predicting the probability of reporting a 2D spatial strategy. Reference levels: Image group, self-reflection, Female.

| Predictor | $\beta$ | OR | SE | p |
|---|---|---|---|---|
| Intercept | $-2.170$ | $0.11$ | $0.435$ | $< .001$ |
| Group (Language vs. Image) | $-0.535$ | $0.59$ | $0.439$ | $.223$ |
| Question (Teaching vs. Self-reflection) | $1.086$ | $2.96$ | $0.433$ | $.012$ |
| Gender (Male vs. Female) | $-0.350$ | $0.70$ | $0.485$ | $.470$ |
| Age ($z$-scored) | $-0.250$ | $0.78$ | $0.249$ | $.316$ |
| $\text{UE}_z$ (Unusual Experiences) | $-0.153$ | $0.86$ | $0.229$ | $.502$ |
| $\text{CD}_z$ (Cognitive Disorganization) | $-0.544$ | $0.58$ | $0.266$ | $.041^*$ |
| $\text{IA}_z$ (Introvertive Anhedonia) | $0.431$ | $1.54$ | $0.218$ | $.048^*$ |
| $\text{IN}_z$ (Impulsive Nonconformity) | $0.351$ | $1.42$ | $0.237$ | $.139$ |

*Note.* OR = odds ratio; SE = standard error. Subscript $z$ denotes $z$-scored predictors. UE, CD, IA, IN = subscales of the Oxford–Liverpool Inventory of Feelings and Experiences (O-LIFE; Mason et al., 1995). The model formula was: logit $P(\text{2D}) = \beta_0 + \beta_1$ group $+ \beta_2$ question $+ \beta_3\,\text{UE}_z + \beta_4\,\text{CD}_z + \beta_5\,\text{IA}_z + \beta_6\,\text{IN}_z + \beta_7\,\text{age}_z + \beta_8$ gender. $^*p < .05$.

Supplementary Table 5 | Tests of the cognitive disorganization × learning-modality interaction and control analyses.

| Test | Estimate | p |
|---|---|---|
| *Strategy reports (categorical outcome)* | | |
| Low vs. high CD, 2D at teaching, image condition (11/30 vs. 3/20) | $OR = 3.28$ | .118 |
| Low vs. high CD, transition to 2D, image condition (10/29 vs. 2/18) | $OR = 4.21$ | .095 |
| Group × CD, teaching reports only | $\beta = 0.381$ | .465 |
| Group × CD, teaching reports, with covariates[a] | $\beta = 0.322$ | .582 |
| Group × CD, transition to 2D | $\beta = 0.125$ | .852 |
| Group × question × CD, all 214 reports | $\beta = 1.014$ | .263 |
| Group × question × CD, proportional-odds model[b] | $\beta = 0.475$ | .405 |
| CD main effect, proportional-odds model[b] | $\beta = -0.126$ | .538 |
| Group difference in transition to 2D (12/47 vs. 4/51)[c] | $\beta = -1.333$ | **.038** |
| *Semantic relevance (continuous outcome)* | | |
| Group × CD, no speech covariates | $b = 0.0074$ | $6.1 \times 10^{-5}$ |
| Group × CD, adding word count | $b = 0.0069$ | $2.1 \times 10^{-4}$ |
| Group × CD, adding word count, sentence count, speech rate, duration | $b = 0.0067$ | $3.8 \times 10^{-5}$ |
| *CD and speech production (Pearson $r$ across reports)* | | |
| Word count (image / language) | $r = -.20$ / $+.12$ | .049 / .202 |
| Sentence count (image / language) | $r = +.01$ / $+.12$ | .887 / .204 |
| Speech duration (image / language) | $r = -.06$ / $-.01$ | .543 / .893 |
| Pause count (image / language) | $r = -.10$ / $+.07$ | .324 / .444 |
| Speech rate (image / language) | $r = -.02$ / $+.10$ | .879 / .290 |
| *Repeated-measures robustness of the logistic model in Table S4*[d] | | |
| Question (teaching), participant-clustered SE | $\beta = 1.086$ | .004 |
| Question (teaching), random-intercept model | $b = 1.727$ | .007 |
| $\text{CD}_z$, participant-clustered SE | $\beta = -0.544$ | .045 |
| $\text{CD}_z$, random-intercept model | $b = -0.974$ | .070 |
| $\text{IA}_z$, participant-clustered SE | $\beta = 0.431$ | .081 |

*Note.* Fisher's exact tests for the first two rows; logistic regression otherwise. CD was $z$-scored. The image condition and self-reflection are the reference levels, so a positive group × CD term indicates a less negative CD slope in the language condition.

[a] Adjusted for the other O-LIFE subscales, age, and gender.

[b] Proportional-odds model with the ordered outcome none < 1D < 2D.

[c] Among participants who did not already report a 2D strategy during self-reflection.

[d] The model of Supplementary Table 4 refitted with standard errors clustered by participant (107 clusters), and as a logistic model with a random intercept per participant (Gauss–Hermite quadrature; $\sigma_u = 2.33$). Coefficients for the remaining predictors did not change in significance under either correction.

Supplementary Table 6 | Excerpts from the verbal reports of two participants in the image condition.

| Participant | Question | Excerpt |
|---|---|---|
| 351 (CD = 4, low) | Self-reflection | "... I tried to make some made-up, like, a story. Usually, like, for breakfast a child should eat, like, a porridge. However, here it was like a teenager was eating, like, a porridge in the morning. And my idea was basically I tried to remember at least four of them, and the fifth one and sixth one I could guess from what is left." |
| 351 | Teaching others | "... there was a diagram about, like, a two-dimensional diagram, where it was like a y-axis was the age of people and x-axis was time. And when I put, like, each food in a specific, like, in a matrix, this actually, like, helped me, like, to better memorize ... Before, what I was using, I was using, like, one-dimensional." |
| 313 (CD = 8, high) | Self-reflection | "... at first, I tried to remember by the appearance ... But then as it got a bit more complex, and I tried to associate it by creating a narrative. So the different appearances, I kind of associated with different people in my life or different time of point in my life." |
| 313 | Teaching others | "... create an image of a restaurant. And the different food items are there, and different people are sitting there ... when you're trying to memorize it's first to associate the food items with the time of the day ... And then to also to connect to the people, the faces, then to think about their relatives that they have." |

*Note.* Both participants learned from images and described no two-dimensional structure during self-reflection. Participant 351 (low CD) introduced an explicit two-dimensional layout when teaching; participant 313 (high CD) described pairwise associations at both questions. Participants were selected as the report closest to the median semantic relevance of their cell among English-language reports, to avoid selection on content; excerpts are verbatim transcriptions, including disfluencies, with omissions marked by ellipses.

### Supplementary results

#### Group comparability

Participants in the image-based and language-based learning conditions did not differ in age, gender, years of education, English proficiency, or language of the verbal report, nor in cognitive disorganization or the other O-LIFE subscales, Big Five personality traits, positive and negative affect, mental rotation, or working memory performance (Supplementary Table 1). The only exception was Impulsive Nonconformity, which was higher in the language-based group; this subscale was included, together with the other O-LIFE subscales, as a covariate in all regression models and showed no association with 2D strategy reports (Supplementary Table 4). Accuracy in the learning-phase tasks (ordering, association learning, navigation) was likewise comparable between groups, indicating that both groups acquired the food–attribute associations to a similar level. The language-based group, however, needed more trials and more time to reach criterion in the ordering and association-learning tasks (Supplementary Table 1). This asymmetry argues against the possibility that the more frequent spatial explanations after image-based learning merely reflect more successful learning in that group.

#### Control analyses of linguistic and acoustic features

To assess whether the observed increase in spatial (2D) strategy reports during teaching could be explained by general differences in language production, we compared text-level and speech-level features of the verbal reports (Supplementary Table 1, lower part). We examined (i) text length (word count, sentence count) and (ii) speech production characteristics (speech duration, pauses, and speech rate). For each feature, we tested (a) differences between the learning conditions within each question, (b) within-condition differences between self-reflection and teaching others, and (c) whether these differences differed between the image-based and language-based learning conditions (i.e., group $\times$ question interaction).

No text-level or speech-level feature showed a significant difference between self-reflection and teaching in either condition, and no feature exhibited a significant group $\times$ question interaction (Supplementary Table 1), indicating that the increase in spatial explanations following visual learning cannot be attributed to differences in response length or speech production. Speech duration was somewhat longer in the image-based than in the language-based group during self-reflection, but this difference was present before the teaching manipulation and did not change differentially across questions. The semantic relevance of the reports to the learned material, quantified with multilingual sentence embeddings, is analyzed in the main text (Figure 3); the sentence-level distribution behind that analysis is shown by question in Supplementary Fig. 4 and, recomputed on English translations of all reports, in Supplementary Fig. 5.

#### Specificity of the cognitive disorganization effect

The logistic regression reported in Supplementary Table 4 is shown as a forest plot in Supplementary Fig. 1. Of the four O-LIFE subscales, only Cognitive Disorganization (CD) was associated with a lower probability of reporting a 2D strategy ($OR = 0.58$, 95

Because the analyses in Figure 2C–D of the main text use a median split, we also plotted the model-implied probability of reporting a 2D strategy as a continuous function of the CD score, with all other covariates held at their sample means (Supplementary Fig. 2). The decline with CD is smooth and the binned observed proportions follow it closely, indicating that the median split is a device for display rather than a driver of the result.

#### Continuous tests of the CD $\times$ learning-modality interaction

The main text reports that the increase in 2D strategy reports from self-reflection to teaching was significant in low-CD participants after image-based learning but not in high-CD participants. Because a difference between a significant and a non-significant within-group test is not itself a test of the difference, we examined the interaction directly, under four specifications (Supplementary Table 5). None supported a CD $\times$ modality interaction on strategy reports: the direct comparison of low- and high-CD participants within the image condition (11/30 vs. 3/20 at teaching, Fisher's exact $p = .118$; 10/29 vs. 2/18 for the transition to 2D, $p = .095$), a logistic model of the teaching reports with a group $\times$ CD term ($\beta = 0.381$, $p = .465$; $\beta = 0.322$, $p = .582$ with covariates), a logistic model of the transition to 2D ($\beta = 0.125$, $p = .852$), and a three-way group $\times$ question $\times$ CD model on all 214 reports ($\beta = 1.014$, $p = .263$). A proportional-odds model treating the outcome as ordered (none < 1D < 2D), which uses more of the information in the ratings, likewise showed no interaction ($\beta = 0.475$, $p = .405$).

Two results are informative rather than merely null. First, the group difference in the teaching effect itself is supported directly: among participants who did not already report a 2D strategy during self-reflection, 12 of 47 in the image condition but only 4 of 51 in the language condition went on to describe 2D structure when teaching ($\beta = -1.333$, $p = .038$). The modality dependence of the teaching effect therefore does not rest on comparing two within-group tests. Second, in the proportional-odds model CD did not shift reports along the ordered scale as a whole ($\beta = -0.126$, $p = .538$) even though it predicted the 2D versus non-2D contrast in the binary model. This pattern is consistent with CD affecting the construction of two-dimensional structure specifically, rather than reducing organized description in general.

We conclude that the modality-specific effect of CD is established by the continuous semantic-relevance analysis (Figure 3), and that the categorical strategy data are consistent with it but individually underpowered: with 22 two-dimensional reports at teaching, the study has limited power to detect an interaction on a binary outcome.

#### Semantic relevance: paired change and control analyses

Semantic relevance decreased from self-reflection to teaching in both conditions, with every participant's pair of reports shown in Supplementary Fig. 3. The relation between semantic relevance and CD is shown separately for each question and condition in Supplementary Fig. 4: it is present for both questions after image-based learning (self-reflection $r = -0.33$, $p = .021$; teaching $r = -0.46$, $p < .001$) and for neither question after language-based learning ($r = 0.16$, $p = .224$ and $r = 0.19$, $p = .160$). As a stronger test of language effects than covariate adjustment, we repeated the analysis on English translations of all 214 reports, re-embedded with the same encoder (Supplementary Fig. 5); the pattern and the group $\times$ CD interaction were unchanged. A concern for the interpretation of Figure 3 is that CD might be associated with how much participants said rather than with what they said. Across reports, CD was essentially unrelated to sentence count, speech duration, pause count, and speech rate in both conditions (all $|r| \leq .12$, all $p > .20$); the only association was a weak negative correlation with word count in the image condition ($r = -.20$,

$p = .049$). The group $\times$ CD interaction on semantic relevance was unchanged when word count was added as a covariate ($b = 0.0069$, $p = 2.1 \times 10^{-4}$) and when word count, sentence count, speech rate, and speech duration were added together ($b = 0.0067$, $p = 3.8 \times 10^{-5}$), compared with $b = 0.0074$, $p = 6.1 \times 10^{-5}$ without speech covariates.

#### Binary analysis of spatial (2D) strategy reports

We additionally analyzed 2D strategy reports as a binary outcome. Exact McNemar tests revealed a significant increase in 2D strategy reports from self-reflection to teaching in the image-based condition (3/50 vs. 14/50; $p = .003$), but not in the language-based condition (6/57 vs. 8/57; $p = .688$; Supplementary Table 3).

Together, these control analyses indicate that the selective increase in spatially structured explanations following visual learning is not driven by general differences in text length or speech production, but instead reflects a specific change in the spatial organization of expressed knowledge.

#### Spatial strategies and cognitive performance

To examine whether spatial strategy use was associated with cognitive performance, we compared behavioral measures between participants who reported a 2D spatial strategy during self-reflection ($n = 9$) and those who did not ($n = 98$), using Wilcoxon rank-sum tests (Supplementary Fig. 6). These analyses were exploratory and were not corrected for multiple comparisons.

Behavioral measures included two-dimensional (2D) placement error, defined as the mean Euclidean distance between each participant's dragged item positions and the true locations in the conceptual space (higher values indicate less accurate spatial memory), corrected N-back accuracy, and mental rotation performance.

Participants who reported 2D spatial strategies during self-reflection showed lower 2D placement error ($M = 0.807$ vs. $M = 1.030$; $p = .007$) and higher corrected N-back accuracy: $M = 1.722$ vs. $M = 1.408$; $p = .010$). No significant association was found with mental rotation performance ($p > .10$). These results suggest that participants who spontaneously used spatial strategies to describe their own memory had better conceptual map accuracy and working memory, consistent with the interpretation that spatial language reflects underlying differences in the quality of conceptual representations.
